# Circulating proteins reveal prior use of menopausal hormonal therapy and increased risk of breast cancer

**DOI:** 10.1101/2021.05.20.444934

**Authors:** Cecilia E. Thomas, Leo Dahl, Sanna Byström, Yan Chen, Mathias Uhlén, Anders Mälarstig, Kamila Czene, Per Hall, Jochen M. Schwenk, Marike Gabrielson

## Abstract

**Background:** Risk prediction is crucial for early detection and prognosis of breast cancer. Circulating plasma proteins could provide a valuable source to increase the validity of risk prediction models, however, no such markers have yet been identified for clinical use.

**Methods:** EDTA plasma samples from 183 breast cancer cases and 366 age-matched controls were collected prior to diagnosis from the Swedish breast cancer cohort KARMA. The samples were profiled on 700 circulating proteins using an exploratory affinity proteomics approach. Linear association analyses were performed on case-control status and a data-driven analysis strategy was applied to cluster the women on their plasma proteome profiles in an unsupervised manner. The resulting clusters were subsequently annotated for the differences in phenotypic characteristics, clinical parameters, and genetic risk.

**Results:** Using the data-driven approach we identified five clusters with distinct proteomic plasma profiles. Women in a particular sub-group (cluster 1) were significantly more likely to have used menopausal hormonal therapy (MHT), more likely to get a breast cancer diagnosis, and were older compared to the remaining clusters. The levels of circulating proteins in cluster 1 were decreased for proteins related to DNA repair and cell replication and increased for proteins related to mammographic density and female tissues. In contrast, classical dichotomous case-control analyses did not reveal any proteins significantly associated with future breast cancer.

**Conclusion:** Using a data-driven approach, we identified a subset of women with circulating proteins associated with previous use of MHT and risk of breast cancer. Our findings point to the potential long-lasting effects of MHT on the circulating proteome even after ending the treatment, and hence provide valuable insights concerning risk predication of breast cancer.

**Highlights:**

- Current risk prediction models use a variety of factors to identify women at risk of developing breast cancer.
- Proteins circulating in blood represent an attractive but currently still underrepresented source of candidates serving as molecular risk factors.
- Plasma proteomes from women participating in a prospective breast cancer cohort study were studied for proteomic risk factors related to a future breast cancer diagnosis.
- Using data-driven approaches, women with future breast cancers and previous use of menopausal hormone therapy were identified based on their circulating proteins.
- Menopausal hormone therapy was found to altered the levels of the circulating proteins even years after the treatment ended.

## Introduction

Breast cancer is the most common cancer among females worldwide and the leading cause of cancer-related mortality in middle aged women [1]. Improving risk prediction and early detection is crucial for better prognosis and survival. Circulating biomarkers have a great potential for simple and minimally-invasive health assessment. Although studies show promising results for blood tests detecting common cancers of the ovary, liver, stomach, pancreas, esophagus, colorectum and lung by circulating proteins [2], identifying putative biomarkers for risk prediction and early detection of breast cancer has thus far been less successful [2-4]. One reason could be that many breast cancers are already being detected at an early stage in mammographic screening programs. Blood levels of early-stage cancer biomarkers are expected to be low and may be too low to detect before the tumor can be uncovered by mammography. Further complicating the search for biomarkers, breast cancer, like most cancers, does not represent a single homogeneous phenotype but consists of multiple subtypes, each arising from particular molecular mechanisms and progressing on distinct clinical paths. So far, proteomic studies have suggested that plasma protein biomarkers for breast cancer may be both subtype and stage specific [3, 5-8]. In addition, there is a growing awareness about inter-individual diversity of molecular profiles even across clinically healthy individuals [9]. Moreover, germline genetic variation may be adding yet another layer of complexity to efforts for finding circulating proteins as common disease biomarkers [10].

Phenotypic and molecular heterogeneity is often limiting the utility of classical dichotomous case-control analyses, as these can prove difficult to delineate or are too simplistic for understanding the underlying molecular subtypes. In these instances, alternative strategies, such as unsupervised and data-driven methods, can allow for novel hypotheses and finding translational biomarkers. Our ambition is to yield unexpected patterns in the data to deliver subgroups that can then readily be linked to molecular phenotypes, clinical risk factors and potentially stratified intervention. Machine learning based clustering is one approach to achieve such explorative, data-driven subtyping and it has been applied successfully in other disease areas such as diabetes [11] and heart failure [12]. Clustering approaches have also previously been applied in breast cancer for prognosis stratification [13, 14] and tumor subtyping [13, 15, 16] using a variety of clinical and molecular parameters. We here used data-driven clustering to stratify women by decomposing their molecular profiles as defined by circulating proteins, and to study the resulting groups for breast cancer risk and risk factors.

With access to the Swedish prospective population-based KARMA cohort [17, 18] we applied exploratory profiling of circulating proteins using a multiplexed affinity proteomics approach based on antibody suspension bead array (SBA) assays. The method allows for many proteins to be screened in small plasma volumes of a large number of samples [19]. To identify proteins associated with phenotypic traits and breast cancer risk factors, we used a data-driven clustering approach and samples from age-matched breast cancer cases and controls collected prior to diagnosis. Our aim was to disentangle the heterogeneity in breast cancer development and risk by improving our limited knowledge about how risk factors influence the plasma proteome and determine if circulating proteins can aid in identifying those individuals at risk of developing breast cancer.

## Material and methods

### Study design, sample inclusion criteria and data collection

The source population was the Karma Cohort consisting of 70,877 women visiting any of four Swedish mammography units during 2011-2013 [17, 18]. All participants signed informed consent forms before joining the KARMA study, and the ethical review board of Karolinska Institutet approved the study. Cases were defined as women diagnosed with breast cancer (N=183) after entering the cohort. Controls were 1:2 matched to each case based on age at last normal screening mammogram and study site (Figure 1).

**Figure 1:**
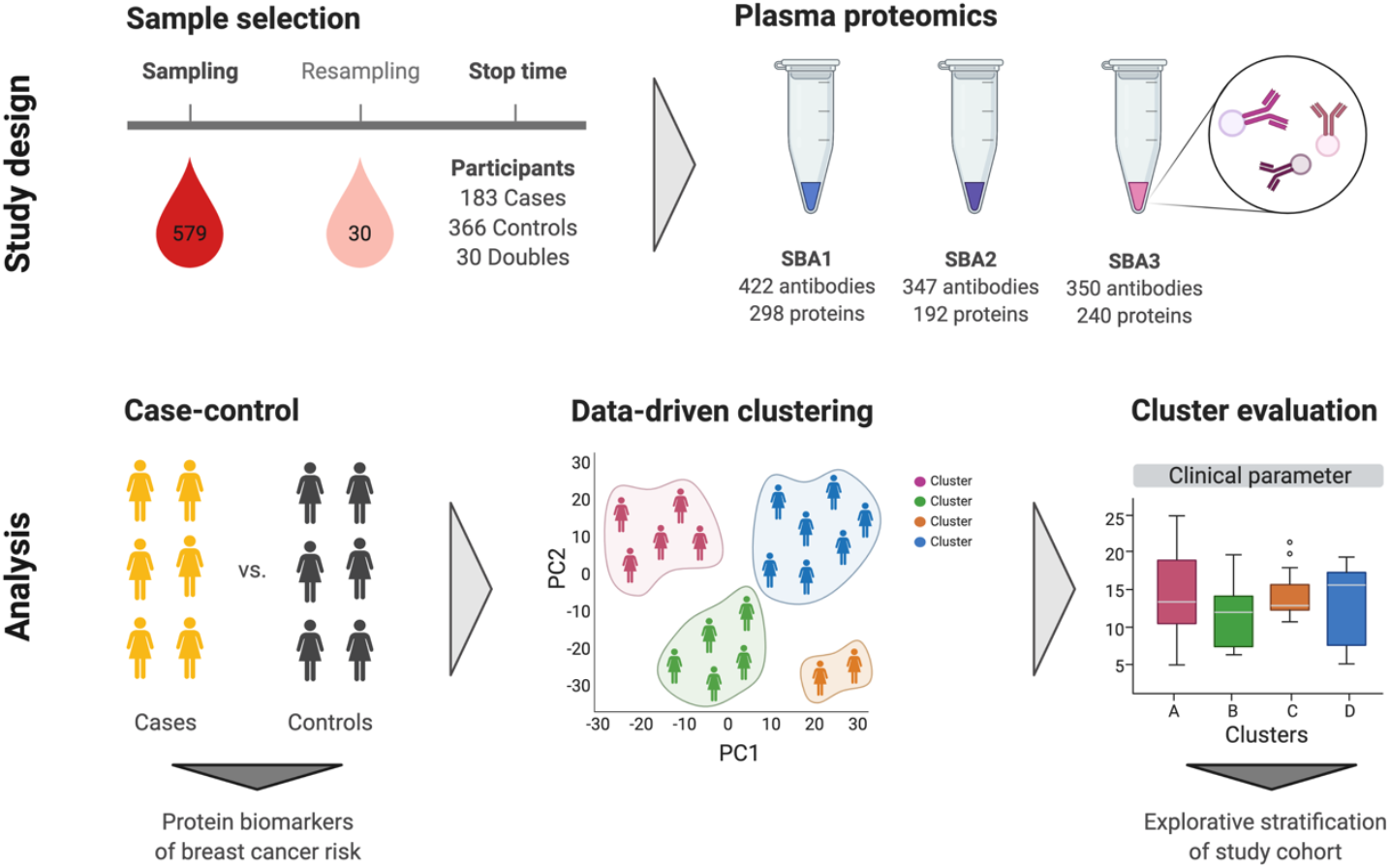
Overview of study design and data analysis. SBA; suspension bead array.

Median time from blood draw to breast cancer diagnosis was 24 days (range 0-588 days). 12 of the incident cases had been diagnosed with breast cancer in the past (5-30 years prior to blood draw; median 11 years). For all, the previous breast tumor was located in the other breast than the tumor that was detected after sampling. 2 controls had previous breast cancer diagnoses, 6 and 16 years prior to study entry. In addition, 19 cases and 10 controls had been diagnosed with other types of cancer prior to sampling (cases: 0.6-35 years; controls: 1-46 years). An additional set of 60 samples from 30 cancer-free individuals from the KARMA cohort were included for quality control (named ‘doubles’). These 30 individuals were sampled on two separate occasions with a median time interval of 19.1 months (range 10.7-19.9) between sampling times.

Raw (unprocessed) digital mammograms for each study participant were collected at KARMA study enrolment as previously described [17, 20]. Additional phenotypic information was obtained from the KARMA study questionnaire and information from national health care registers [17]. BMI was calculated at time of study entry and was based on self-reported height and weight. Information on tumor characteristics was obtained by linkage to the Swedish nation-wide cancer registry. Information on menopausal hormonal therapy (MHT) and statin use was extracted from the Swedish drug prescription registry. Anatomical Therapeutic Chemical (ATC) codes were extracted for MHT containing only estrogens, only progestogens or a combination of estrogens and progestogens, as well as for lipophilic and hydrophilic statins (**Supplementary table S1**).

### Plasma sample collection

Non-fasting EDTA plasma samples of peripheral blood were collected from the KARMA study participants at enrolment [17]. All blood samples were handled in accordance with a strict 30-hours cold-chain protocol and were processed in the Karolinska Institutet high-throughput biobank. Samples were collected between January 2011 and September 2012.

### Antibody bead arrays

We used antibody suspension bead arrays (SBA) to determine protein profiles in plasma samples. The SBAs were generated using carboxylated magnetic beads (MagPlex-C, Luminex Corp.) as previously described [19]. All plasma samples within each study set were retrieved from the biobank and analyzed at the same point in time. Plasma samples stored at - 80°C were thawed at 4°C and randomized across seven 96-well microtiter plates in a stratified manner: Each double pair and trio (case and two matched controls) were placed within the same plate, resulting in an even distribution of cases, controls and doubles across all seven plates. Samples were assayed in 384 well plates, where the fourth quadrant in each 384-well contained the same 96 samples that originated from one of the crude 96-well sample plates. In addition, all plates included four aliquot replicates from a crude plasma pool from all individuals included in the study. Samples were biotinylated, diluted, heat-treated at 56°C and combined with the bead array on two separate 384-well assay plates in accordance with previously described protocols [21]. The protein levels were reported as units of the median fluorescence intensity (MFI) from measuring at least 32 beads per antibody assay.

### Protein target selection

We used antibodies derived from the Human Protein Atlas [22] to construct three SBAs were built on sets of 422, 347 and 350 antibodies (SBA1-SBA3, **Supplementary Figure S1**) as previously described [9]. These targeted a total 729 unique protein-encoding genes, and a complete list of all antibodies included in the study is provided in **Additional file 1**. The 422 antibodies included in the first bead array (SBA1) targeted 295 protein-encoding genes annotated to extracellular matrix [Uniprot.org] [23], including integrins (N=27), laminins (N=21), matrix metalloproteases (N=21), metallopeptidases (N=18), and proteoglycans (N=16). A majority of the antibodies (82%) in SBA1 targeted secreted proteins. The 347 antibodies in SBA2 included 243 antibodies (127 proteins) targeting breast cancer-related proteins from literature, 62 antibodies towards 55 proteins with strong expression in breast tissue according to RNAseq data [proteinatlas.org], 39 antibodies towards 11 proteins with indicative associations to breast cancer from previous screenings and 3 controls. The 350 antibodies against 241 protein-encoding genes included in the third suspension bead array (SBA3) were selected based on possible relationship to mammographic breast density, cancer development and/or progression, or tissue composition and/or remodeling. Due to overlap between the different arrays the total number was 1,073 unique antibodies targeting 701 unique proteins. This included sets of paired antibodies with common protein targets.

### Data processing

The generated raw protein profile data was normalized and annotated as follows. Antibody-specific probabilistic quotient normalization (Abs-PQN) [9] was applied per 96-well plate to reduce within-plate sample-to-sample variation. Between-plate normalization was performed using a multidimensional (MA) normalization method [24] (Supplementary Figure S2).

A set of 96 duplicated samples were used to assess technical variation and to confirm reproducibility of antibody profiles within all three SBAs. Prior to statistical analyses, the data were annotated based on assay performance using three criteria. Internal controls and antibodies were excluded from proceeding analyses if they showed low reproducibility in replicated analyses (as rho<0.7), correlation to human IgG levels (rho>0.5), or elevated background levels in assays with sample-free buffers (MFI_Empty_ > mean(MFI_Sample_) + 3×sd(MFI_Sample_)). Replicates samples were also excluded prior to the analyses.

### Case-control analysis

For contrasting cases versus controls, conditional logistic regression models considering the age- and sampling location matching of cases and controls were applied to normalized, Ab-filtered and log transformed data. Three models were compared. In model 1, BMI and study entry date were included as exposure variables. Model 2 included exposure variables for absolute area-based breast density, postmenopausal status (yes/no) and MHT use (yes/no) in addition to BMI and entry date. In model 3, smoking (packs/year), alcohol (grams/week) and childbirth (yes/no) were included as exposure variables in addition to the variables in model 2. Due to missing values for BMI (4 missing), area-based density (20 missing), MHT usage (5 missing), smoking (3 missing), alcohol (2 missing), and childbirth (1 missing), 540 samples (181 cases, 359 controls) were analyzed in model 1, 490 samples (167 cases, 323 controls) were analyzed in model 2 and 484 (165 cases, 319 controls) were analyzed in model 3. Statistical modeling was performed using the “clogit” function of the “survival” R package (version 3.1.8) [25, 26].

### Unsupervised clustering

We performed an unsupervised archetype clustering of the proteomics data to identify clusters of individuals with similar protein profiles. These profiles were subsequently associated to clinical risk factors and other traits.

The quality-controlled proteomics data was linearly adjusted for BMI, entry date, and age at sampling. Clustering was performed using archetypal analysis where each individual can be described as a combination of archetypes that represent extremes in the data. Archetypal analysis was performed using the “archetypal” function of the “archetypal” R package (version 1.1.0) [27]. After archetypal analysis clusters were created by assigning each individual to the archetype that they had the highest probability of belonging to. To validate the clusters we tested the stability of the clusters when the data was changed slightly [28]. This was done by bootstrap analysis where a subset of patients was randomly selected, and the clustering performed on the subset and the results compared to the clustering on the original data. For technical assessment of the clustering, the results of the archetypal analysis were used to predict the archetype coefficients of doubles and replicates that had been excluded from the original clustering. This was done using the “predict” function of the “stats” R package (version 3.6.0) on an “archetypes” object of the “archetypes” R package (version 2.2.0.1) [29]. Further details on the clustering analysis can be found in the supplementary material.

### Statistical tests of cluster characteristics

We compared the clusters to investigate how the differences in protein levels driving the clustering materialized at the clinical level. Similarly, we compared the genetic predisposition to breast cancer to assess if the differences in protein levels might be genetically driven. Details on the genetic data and calculation of polygenic risk scores (PRSs) are given in the supplementary material. The Wilcoxon rank-sum test was used for continuous variables and Fisher’s exact test for categorical variables. Testing of the influence of potential genetic components between the clusters was done by the absolute values of PRS in the clusters as a continuous variable. All *P-*values were two sided and considered statistically significant if <0.05.

To rank the proteins driving a cluster, we first performed differential abundance analysis comparing a cluster to the remaining samples using the t-test. Resulting p-values were corrected for multiple comparisons using Benjamini-Hochberg adjustment resulting in false discovery rates (FDRs) for each protein. Next, we performed pathway analysis to summarize the potential functions of differentially abundant circulating proteins. We began by applying Over-Representation Analysis (ORA) using two separate criteria for protein selection; proteins with a FDR < 0.05 and the top 50 proteins with the lowest p-value, using the “gost” function of the “gprofiler2” R package (version 0.1.8) [30]. Next, we applied Gene Set Enrichment analysis (GSEA) where all proteins were included but ranked by their p-value and direction of differential abundance, using the “fgsea” R package (version 1.12.0) from Bioconductor [31].

To shortlist representative proteins for a cluster, we selected the union of those with the lowest p-values and the highest (positive or negative) difference in relative abundance. The levels of the selected proteins in all participants were associated with dense area (adjusted for BMI and age) and MHT status (never taken, taken before study entry, taking at entry) using linear and logistic regression, respectively. All data handling and statistical analyses were performed in R version 3.6.0.

## Results

### Characterizing the cohort

The selected study population consisted of 183 cases and 366 matched controls (**Table 1**), as well as 30 doubles that were sampled twice over time (**Supplementary Table S2**). Cases and controls had similar BMI, but cases had a higher absolute area-based breast density (p = 0.0045). 74.9% of cases were postmenopausal, with similar proportions for controls. 48.1% of cases and 46.7% of controls had never taken MHT, with similar numbers for statin use. The majority of the tumors were positive for ER (74.9%) and PR (59.6%), only a few confirmed HER2 positive (7.7%). More than half of the tumors were invasive (54.1%) with histological grade ≥2 (76.5%) but without lymph node invasion (78.1%). Women were recruited at four centers, but no differences between sampling centers were observed at the protein level (Supplementary Figure S3).

**Table 1:** Overview of clinical characteristics for cases and controls, and tumor characteristics for cases. P-values are from comparing cases and controls using Wilcoxon rank-sum tests for continuous variables and Fisher’s exact tests for categorical variables.

|  | Total<br>(N=549) | Cases<br>(N=183) | Controls<br>(N=366) | P-value |
| --- | --- | --- | --- | --- |
| Age |  |  |  |  |
| Mean (SD) | 59.6 (9.28) | 59.6 (9.30) | 59.6 (9.28) | 1 |
| Median<br>[Min, Max] | 62.0 [39.0, 81.0] | 62.0 [39.0, 81.0] | 62.0 [39.0, 81.0] |  |
| BMI |  |  |  |  |
| Mean (SD) | 25.6 (4.19) | 25.8 (3.78) | 25.5 (4.38) | 0.13 |
| Median<br>[Min, Max] | 24.9 [17.6, 49.0] | 25.4 [18.5, 39.2] | 24.7 [17.6, 49.0] |  |
| Missing | 4 (0.7%) | 1 (0.5%) | 3 (0.8%) |  |
| Sampling center |  |  |  |  |
| Helsingborg Hospital | 283 (51.5%) | 95 (51.9%) | 188 (51.4%) | 0.99 |
| Landskrona Hospital | 23 (4.2%) | 7 (3.8%) | 16 (4.4%) |  |
| Skåne University Hospital, Lund | 20 (3.6%) | 7 (3.8%) | 13 (3.6%) |  |
| Stockholm South General Hospital | 223 (40.6%) | 74 (40.4%) | 149 (40.7%) |  |
| Menopausal status |  |  |  |  |
| Premenopausal | 130 (23.7%) | 45 (24.6%) | 85 (23.2%) | 0.75 |
| Postmenopausal | 418 (76.1%) | 137 (74.9%) | 281 (76.8%) |  |
| Missing | 1 (0.2%) | 1 (0.5%) | 0 (0%) |  |
| Dense area (cm2) |  |  |  |  |
| Mean (SD) | 27.3 (24.2) | 30.9 (24.1) | 25.6 (24.1) | 0.005 |
| Median [Min, Max] | 20.4 [0.0, 161.4] | 23.6 [0.1, 113.6] | 18.7 [0.0, 161.4] |  |
| Missing | 20 (3.6%) | 14 (7.7%) | 6 (1.6%) |  |
| **MHT status** |  |  |  |  |
| Never taken | 259 (47.2%) | 88 (48.1%) | 171 (46.7%) | 0.51 |
| Taken before | 213 (38.8%) | 74 (40.4%) | 139 (38.0%) |  |
| Taking at sampling | 70 (12.8%) | 19 (10.4%) | 51 (13.9%) |  |
| Missing | 7 (1.3%) | 2 (1.1%) | 5 (1.4%) |  |
| **Statin status** |  |  |  |  |
| Never taken | 272 (49.5%) | 86 (47.0%) | 186 (50.8%) | 0.76 |
| Taken before | 47 (8.6%) | 15 (8.2%) | 32 (8.7%) |  |
| Taking at sampling | 52 (9.5%) | 19 (10.4%) | 33 (9.0%) |  |
| Missing | 178 (32.4%) | 63 (34.4%) | 115 (31.4%) |  |
| **Smoking**<br>**(packs per year)** |  |  |  |  |
| Mean (SD) | 6.08 (9.57) | 6.46 (9.73) | 5.89 (9.50) | 0.30 |
| Median [Min, Max] | 0.950 [0, 64.2] | 1.65 [0, 49.3] | 0.800 [0, 64.2] |  |
| Missing | 3 (0.5%) | 3 (1.6%) | 0 (0%) |  |
| **Alcohol intake**<br>**(g per week)** |  |  |  |  |
| Mean (SD) | 58.2 (69.9) | 60.0 (70.9) | 57.3 (69.5) | 0.88 |
| Median [Min, Max] | 37.0 [0, 575] | 37.0 [0, 292] | 37.0 [0, 575] |  |
| Missing | 2 (0.4%) | 2 (1.1%) | 0 (0%) |  |
| **Ever given birth** |  |  |  |  |
| Never given birth | 78 (14.2%) | 27 (14.8%) | 51 (13.9%) | 0.80 |
| Has given birth | 470 (85.6%) | 155 (84.7%) | 315 (86.1%) |  |
| Missing | 1 (0.2%) | 1 (0.5%) | 0 (0%) |  |
| **ER status** |  |  |  |  |
| Negative | - | 18 (9.8%) | - |  |
| Positive | - | 137 (74.9%) | - |  |
| Missing | - | 28 (15.3%) | - |  |
| **PR status** |  |  |  |  |
| Negative | - | 44 (24.0%) | - |  |
| Positive | - | 109 (59.6%) | - |  |
| Missing | - | 30 (16.4%) | - |  |
| **HER2 status** |  |  |  |  |
| Negative | - | 136 (74.3%) | - |  |
| Positive | - | 14 (7.7%) | - |  |
| Missing | - | 33 (18.0%) | - |  |
| **Invasiveness** |  |  |  |  |
| Invasive | - | 99 (54.1%) | - |  |
| Carcinoma in situ | - | 19 (10.4%) | - |  |
| Missing | - | 65 (35.5%) | - |  |
| **Tumor size** |  |  |  |  |
| < 20 mm | - | 43 (23.5%) | - |  |
| >= 20 mm | - | 17 (9.3%) | - |  |
| Missing | - | 123 (67.2%) | - |  |
| **Lymph node metastasis** |  |  |  |  |

|  | **Total**<br>**(N=549)** | **Cases**<br>**(N=183)** | **Controls**<br>**(N=366)** | **P-value** |
| --- | --- | --- | --- | --- |
| No | - | 143 (78.1%) | - |  |
| Yes | - | 15 (8.2%) | - |  |
| Missing | - | 25 (13.7%) | - |  |
| **Nottingham Histologic Grade** |  |  |  |  |
| 1 | - | 31 (16.9%) | - |  |
| 2 | - | 68 (37.2%) | - |  |
| 3 | - | 72 (39.3%) | - |  |
| Missing | - | 12 (6.6%) | - |  |

Abbreviations: Body mass index (BMI), Menopausal hormone therapy (MHT), Estrogen receptor (ER), Progesterone receptor (PR), Human epidermal growth factor receptor 2 (HER2).

### Identifying protein biomarkers of case-control status

A set of 54 proteins were associated with case-controls status with a nominal p < 0.05 in at least one of the three conditional logistic regression models tested (data not shown). However, after adjustment for multiple testing none remained significant (FDR > 0.05).

### Unsupervised clustering of participants based on their protein profiles

Prior to clustering we adjusted the proteomics data for a selected set of covariates. The impact of BMI, age of the women at sampling and study entry date (as a proxy for sample age) on the protein data were studied by projecting the data to two dimensions using Uniform Manifold Approximation and Projection (UMAP) (**Supplementary Figure S4**) and by associating protein levels with BMI, age and entry date in a combined linear model. The linear association resulted in significant (p < 0.05) associations for 305, 415 and 57 proteins for BMI, age and entry date respectively. Thus, when considering both the overall impact on the measured proteins and the effect on individual proteins, age of the women had the strongest influence on the measured proteins as a whole, followed by BMI and with a limited effect of entry date. The experimental proteomics data were therefore adjusted for BMI, age of the women, and study entry date prior to further analyses. Five individuals lacked information on BMI and were therefore excluded, leaving 573 samples (181 cases, 363 controls, 29 doubles) for analysis. 552 unique antibodies with 552 unique targets were left after removing antibodies with the same target.

To identify patterns in the proteomics data grouping individuals into clusters, we performed archetypal analysis. We applied the Unit Invariant Knee method to identify the optimal number of clusters (as described in the supplementary material) (**Supplementary Figure S5**) that would balance simplicity with adequate stratification of the data. This resulted in 5 clusters with 19, 113, 115, 144, and 182 participants respectively (**Figure 2A-2D**), representing 3.32%, 19.7%, 20.0%, 25.1% and 31.8% of all tested subjects.

**Figure 2:**
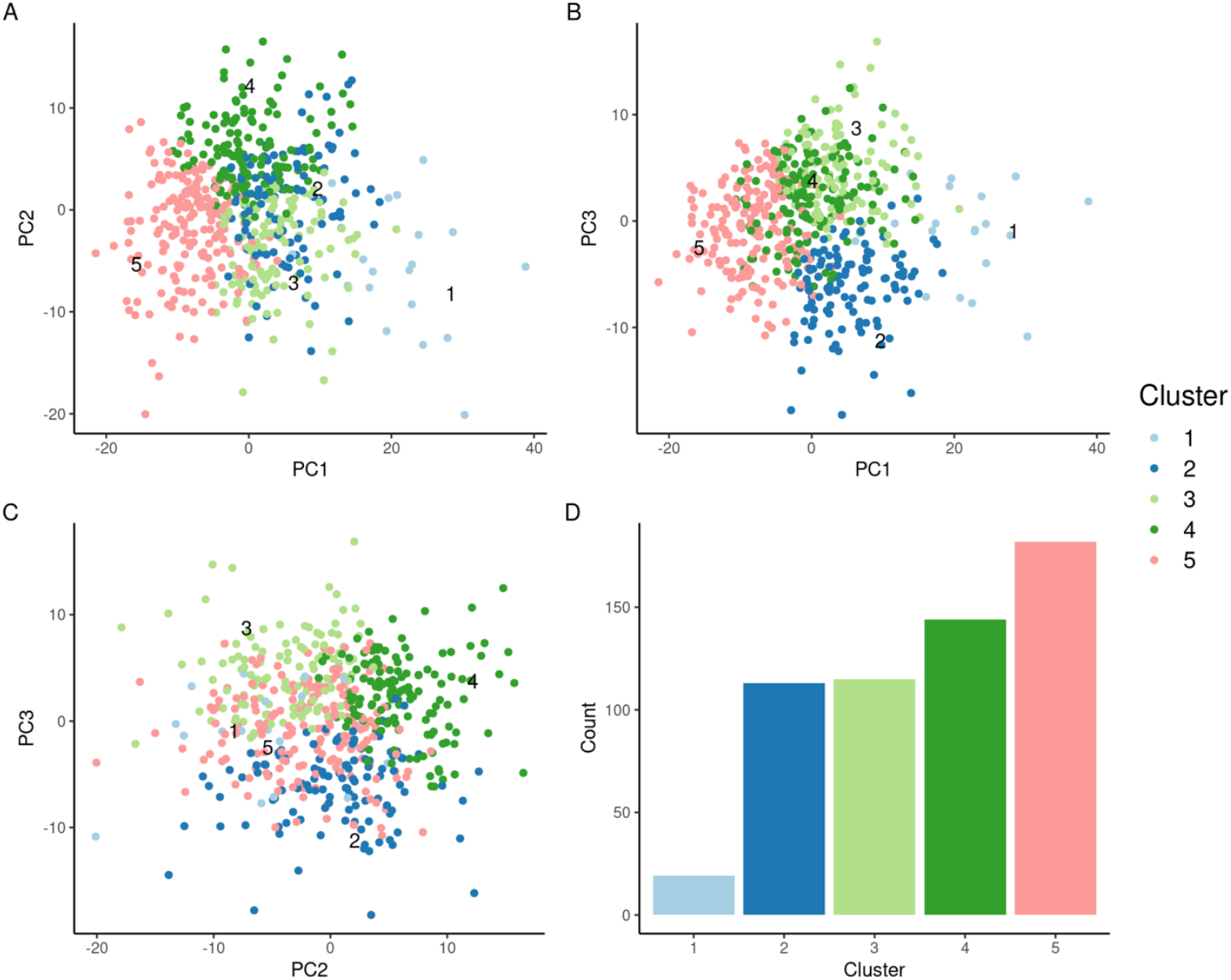
Principal component analysis (PCA) of each participant’s protein profile plotted with (A) PC1/PC2, (B) PC1/PC3, and (C) PC2/PC3. Each dot represents one participant. Participants are colored by which cluster they were assigned to. (D) number of participants in each cluster.

The mean Jaccard index, used to assess the cluster stability, was 0.48, 0.35, 0.34, 0.37, and 0.36 for cluster 1-5, respectively (**Supplementary Table S3**). To further assess the quality of the clustering, we determined the cluster membership of pairs of replicated samples and pairs of samples collected on different occasions from the same individual (double samples). We observed that replicate sample pairs significantly more often belonged to the same cluster than double sample pairs (**Supplementary Figures S6-S8, supplementary results**). This is in line with the difference in measured protein levels between replicate sample pairs being of purely technical origin, while differences in measured protein levels of the double pairs can be of both technical and biological origin due to the time elapsed between samplings. In addition, doubles pairs belonged more often to the same cluster than random pairs of samples. Thus, also showing that the protein profiles of the individual women did not substantially change between samplings. Taken together, this indicated that the clustering captures groups of individuals with similar protein profiles.

### Clinically characterizing the clusters of participants

Clusters of participants were defined at the protein level, and we proceeded to investigate how the stratification observed at the protein level might be reflected at the clinical level. We therefore contrasted a range of clinical variables across the clusters (**Table 2 and Supplementary Table S4**). Women belonging to cluster 1 had distinct clinical characteristics. Given that cluster 1 was the most stable cluster as determined by the Jaccard index and was the cluster with the most unique protein profile, we focused the remaining part of the analyses on this cluster. Cluster 1 consisted of women of higher age compared to clusters 2 and 4 (p < 0.05, **Figure 3A**), despite the proteomics data being adjusted for age prior to archetype clustering. BMI and BMI-adjusted area-based breast density was not significantly different across clusters (**Figure 3B-3C**). Cluster 1 had a mean and median dense area of 25.8 cm^2^ and 21.2 cm^2^, respectively (**Table 2**). Though the density for women in cluster 1was not significantly different than the other clusters, it was substantially higher than a comparative sub-group of women of the same age. The group used for comparison were women within the same age range (63-65) and proportion of breast cancer cases from the KARMA cohort [32, 33].

**Table 2:** Overview of the clinical characteristics of the archetype clusters.

|  | 1<br>(N=19) | 2<br>(N=113) | 3<br>(N=115) | 4<br>(N=144) | 5<br>(N=182) |
| --- | --- | --- | --- | --- | --- |
| **Case control status** |  |  |  |  |  |
| Case | 11 (57.9%) | 32 (28.3%) | 37 (32.2%) | 38 (26.4%) | 63 (34.6%) |
| Control | 8 (42.1%) | 81 (71.7%) | 78 (67.8%) | 106 (73.6%) | 119 (65.4%) |
| **Age** |  |  |  |  |  |
| Mean (SD) | 63.7 (6.95) | 58.7 (9.29) | 59.7 (9.63) | 58.5 (9.97) | 59.1 (9.33) |
| Median | 65.0 | 61.0 | 63.0 | 61.5 | 62.0 |
| [Min, Max] | [46.0, 76.0] | [40.0, 78.0] | [39.0, 81.0] | [40.0, 78.0] | [39.0, 81.0] |
| **BMI** |  |  |  |  |  |
| Mean (SD) | 24.2 (4.10) | 25.5 (3.96) | 25.6 (3.70) | 25.6 (4.65) | 25.3 (4.24) |
| Median | 23.7 | 24.8 | 25.2 | 24.8 | 25.0 |
| [Min, Max] | [17.9, 33.9] | [18.8, 37.0] | [18.5, 36.3] | [18.4, 44.2] | [17.6, 49.0] |
| **MHT status** |  |  |  |  |  |
| Never taken | 4 (21.1%) | 54 (47.8%) | 56 (48.7%) | 68 (47.2%) | 89 (48.9%) |
| Taken before | 14 (73.7%) | 45 (39.8%) | 42 (36.5%) | 59 (41.0%) | 64 (35.2%) |
| Taking at entry | 1 (5.3%) | 14 (12.4%) | 15 (13.0%) | 15 (10.4%) | 27 (14.8%) |
| Missing | 0 (0%) | 0 (0%) | 2 (1.7%) | 2 (1.4%) | 2 (1.1%) |
| **Statin status** |  |  |  |  |  |
| Never taken | 12 (63.2%) | 62 (54.9%) | 55 (47.8%) | 68 (47.2%) | 93 (51.1%) |
| Taken before | 1 (5.3%) | 9 (8.0%) | 8 (7.0%) | 13 (9.0%) | 17 (9.3%) |
| Taking at entry | 2 (10.5%) | 14 (12.4%) | 9 (7.8%) | 8 (5.6%) | 19 (10.4%) |
| Missing | 4 (21.1%) | 28 (24.8%) | 43 (37.4%) | 55 (38.2%) | 53 (29.1%) |
| **Menopausal status** |  |  |  |  |  |
| Premenopausal | 1 (5.3%) | 33 (29.2%) | 27 (23.5%) | 41 (28.5%) | 46 (25.3%) |
| Postmenopausal | 18 (94.7%) | 80 (70.8%) | 88 (76.5%) | 103 (71.5%) | 136 (74.7%) |
| **Dense area (cm2)** |  |  |  |  |  |
| Mean (SD) | 25.8 (20.7) | 29.0 (27.2) | 28.6 (28.7) | 30.0 (26.0) | 25.6 (20.0) |
| Median<br>[Min, Max] | 21.2<br>[1.3, 73.7] | 23.6<br>[0.0, 124.0] | 19.8<br>[0.0, 161.0] | 21.0<br>[0.0, 119.0] | 20.4<br>[0.0, 86.9] |
| Missing | 0 (0%) | 10 (8.8%) | 0 (0%) | 3 (2.1%) | 7 (3.8%) |
| **BMI- and age-adjusted dense area (cm2)** |  |  |  |  |  |
| Mean (SD) | 20.7 (18.3) | 21.6 (24.7) | 22.7 (26.6) | 23.0 (22.0) | 18.7 (18.4) |
| Median<br>[Min, Max] | 15.5<br>[-4.4, 70.3] | 12.7<br>[-12.7, 109.0] | 16.4<br>[-12.6, 161.0] | 16.5<br>[-9.4, 90.2] | 14.8<br>[-13.3, 76.1] |
| Missing | 0 (0%) | 10 (8.8%) | 0 (0%) | 3 (2.1%) | 7 (3.8%) |
| **Smoking (packs per year)** |  |  |  |  |  |
| Mean (SD) | 7.34 (9.69) | 7.08 (10.0) | 6.24 (10.7) | 5.68 (8.01) | 5.21 (9.33) |
| Median<br>[Min, Max] | 1.50<br>[0, 29.1] | 1.50<br>[0, 46.6] | 0<br>[0, 49.3] | 1.50<br>[0, 42.9] | 0.450<br>[0, 64.2] |
| Missing | 0 (0%) | 1 (0.9%) | 1 (0.9%) | 0 (0%) | 0 (0%) |
| **Alcohol intake**<br>**(g per week)** |  |  |  |  |  |
| Mean (SD) | 70.4 (69.6) | 49.2 (64.4) | 52.4 (60.1) | 76.9 (76.4) | 51.3 (70.2) |
| Median<br>[Min, Max] | 37.0<br>[0, 261] | 37.0<br>[0, 362] | 37.0<br>[0, 273] | 37.0<br>[0, 292] | 37.0<br>[0, 575] |
| Missing | 0 (0%) | 0 (0%) | 1 (0.9%) | 0 (0%) | 0 (0%) |
| **Ever given birth** |  |  |  |  |  |
| Never given birth | 5 (26.3%) | 16 (14.2%) | 17 (14.8%) | 13 (9.0%) | 30 (16.5%) |
| Has given birth | 14 (73.7%) | 97 (85.8%) | 98 (85.2%) | 131 (91.0%) | 152 (83.5%) |

Abbreviations: Body mass index (BMI), Menopausal hormone therapy (MHT).

**Figure 3:**
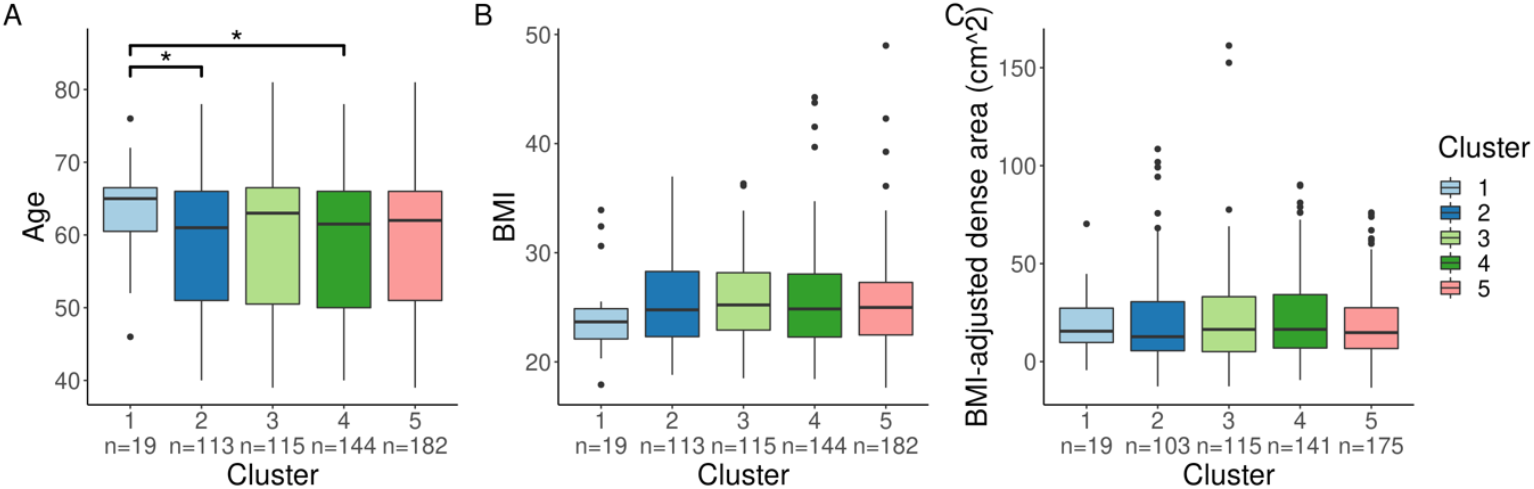
Distributions of (A) age, (B) BMI, and (C) mammographic density (cm^2^) (age- and BMI-adjusted) in the clusters. An asterisk symbolizes a Wilcoxon rank-sum test p < 0.05 by pairwise comparison.

There was a significantly greater proportion of breast cancer cases in cluster 1 compared to the clusters 2, 3, and 4 (all p < 0.05, **Figure 4A**). Cluster 1 also had a significantly greater proportion of women who had taken MHT compared to the other clusters (all p < 0.05, **Figure 4B**). Additionally, the proportion of women who had previously taken MHT prior to study entry but were not taking MHT at the time of blood sampling, was also significantly higher in cluster 1 (all p < 0.05, **Figure 4C**). We observed no significant difference between clusters regarding the time from last MHT to study entry (**Figure 4D**). Cluster 1 contained a higher proportion of cases who had taken MHT ever (100% of cases) compared to other clusters (approximately 50% of cases) (**Figure 4E**).

**Figure 4:**
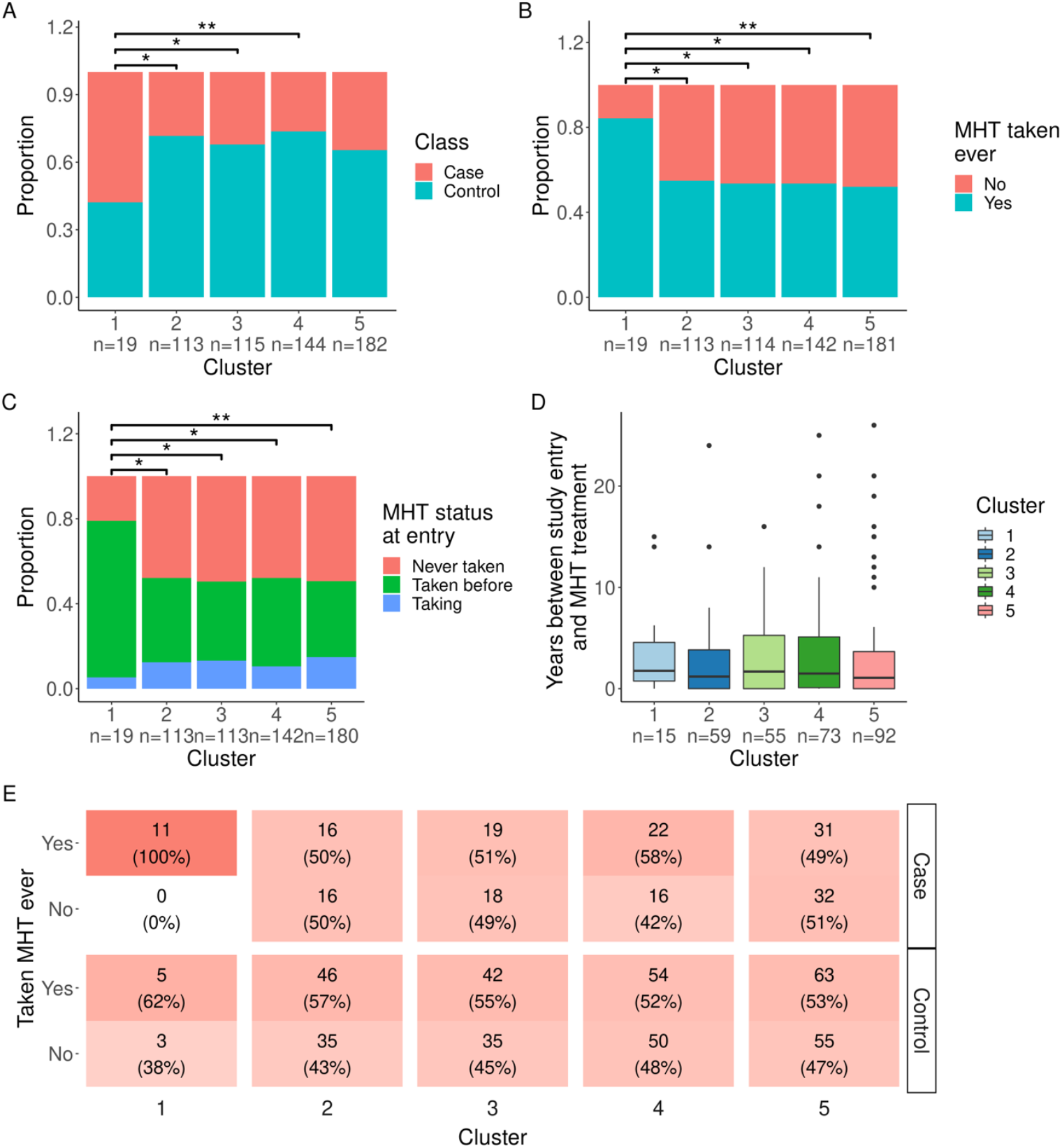
Comparison between the five clusters in proportions of (A) cases and controls, where doubles were treated as controls as they were all cancer-free at study entry. (B) Participants who had ever taken MHT, and (C) participants who were taking MHT prior to sample collection, at time of sampling, or never. (D) Time in years between last use of MHT and study entry for the five clusters. Asterisks symbolize Fisher’s exact test p-values (*: p<0.05, **: p<0.01) for pairwise comparisons between clusters. (E) Numbers of participants in each cluster who have taken or not taken MHT, divided by case-control status.

Given that 101 of the women were currently using or had previously been treated with statins and that statin use has previously been shown to affect the plasma proteome [34, 35], we wanted to exclude this as a possible confounding factor. We observed no significant difference between clusters regarding statin usage, neither when delineating by statin type nor when grouping all statins together (**Supplementary Figure S9**). Lastly, we compared PRSs across clusters and found no significant difference. Also, no significant differences were observed when comparing only cases in cluster 1 with cases in the remaining clusters. Additionally, when comparing the PRS of all case to all controls, the PRS was slightly higher for cases, however, this difference was not statistically significant. This could be due to small sample sizes. (**Supplementary Figure S10**).

Given that several cases and controls had previous cancer diagnoses we reran the clinical comparison of the clusters where these individuals were excluded to ascertain that such previous cancer and related treatment was not driving the differences observed. We did not observe any major changes resulting from excluding these individuals (data not shown).

### Investigating the proteomic differences between clusters of participants

Differences in protein levels between the clusters were observed with a heatmap (**Figure 5A**). Distinct patterns reflecting the differences in protein levels can be observed for all clusters but are most apparent for cluster 1. The differential abundance analysis comparing the protein profiles of women in cluster 1 with all other individuals yielded 393 (72% of all) proteins with higher levels, of which 245 had an FDR < 0.05. In contrast, there were 159 (28% of all) proteins with lower levels, 73 of which had an FDR < 0.05. There were no significantly enriched pathways neither from the ORA over-representation analysis nor the GSEA gene set enrichment analysis. However, this investigation was likely biased by the already highly selective design to target only a particular set of proteins in the circulation.

**Figure 5:**
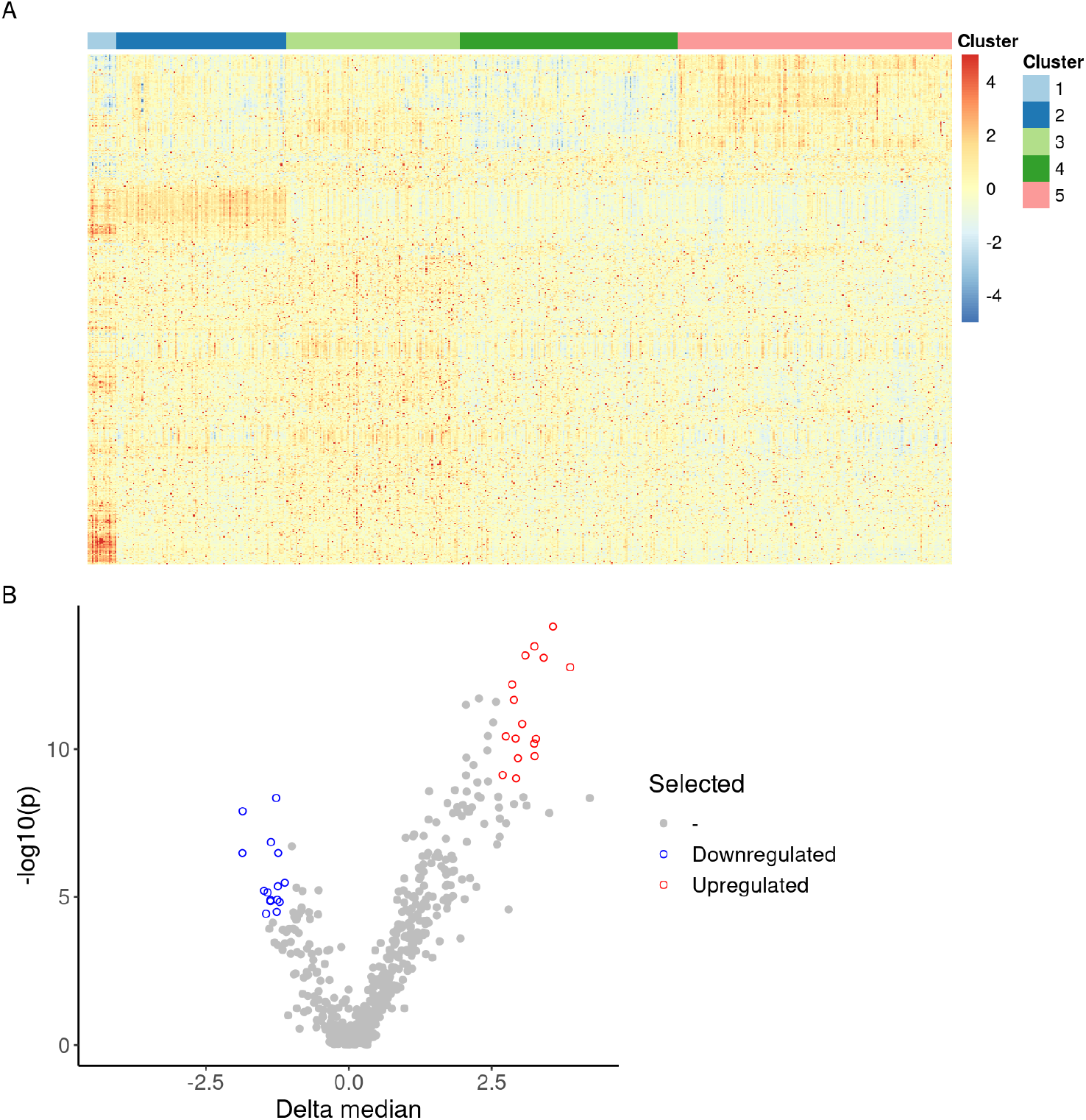
Proteomic characterization of clusters. (A) Heatmap of normalized, centered and scaled MFI for each protein (rows) and participant (columns). Participants are ordered into the archetype clusters they belong to, while proteins are clustered using hierarchical clustering based on Euclidean distance. (B) Volcano plot of differentially abundant proteins in cluster 1 compared to samples in the remaining clusters. Blue: A subset of 14 proteins with lower plasma levels were selected form the union of the 25 proteins with the lowest p-values and the 25 proteins with the largest decrease in abundance levels. Red: A subset of 16 proteins with higher plasma levels selected from the union of the 25 of proteins with the lowest p-values and the 25 proteins with the largest increase in abundance levels.

To provide insights into the proteomic signatures of cluster 1, we shortlisted those proteins unifying the lowest p-values and largest relative abundance differences. Compared to the rest of the participants and choosing the union of the 25 most significant and 25 most differentially abundant proteins of cluster 1 (**Figure 5B**), there were 16 more abundant (**Table 3**) and 15 less abundant proteins (**Table 4**). The levels of PTCH1 and ZP4 were significantly associated with adjusted breast density (nominal p < 0.05) and MHT status (nominal p < 0.05) when performing linear regression and logistic regression, respectively. CCR7, MMRN1, HNRNPA2B1, RBBP8, ACOX3, TJP3, and MMP15 were associated with adjusted breast density (nominal p < 0.05), but not MTH status (**Supplementary Figure S11**). MFI levels of PTCH1 and ZP4 were lower in cases than in controls and significantly lower if MHT had been used (**Supplementary Figure S12**).

**Table 3.** Proteins with lower plasma levels in cluster 1 compared to the other clusters.

| Gene name | Gene description | ENSG ID | FDR | FC |
| --- | --- | --- | --- | --- |
| F11R | F11 receptor | ENSG00000158769 | 1,93E-11 | 3,88 |
| CLDN15 | claudin 15 | ENSG00000106404 | 4,08E-12 | 3,57 |
| EXOC2 | exocyst complex component 2 | ENSG00000112685 | 1,14E-11 | 3,41 |
| CYBB | cytochrome b-245 beta chain | ENSG00000165168 | 1,56E-09 | 3,28 |
| NTN4 | netrin 4 | ENSG00000074527 | 5,14E-09 | 3,26 |
| RNASE2 | ribonuclease A family member 2 | ENSG00000169385 | 9,52E-12 | 3,25 |
| CCR10 | C-C motif chemokine receptor 10 | ENSG00000184451 | 2,14E-09 | 3,25 |
| MET | MET proto-oncogene, receptor tyrosine kinase | ENSG00000105976 | 1,14E-11 | 3,09 |
| MLH3 | mutL homolog 3 | ENSG00000119684 | 6,56E-10 | 3,04 |
| ITGB7 | integrin subunit beta 7 | ENSG00000139626 | 5,4E-09 | 2,96 |
| TIE1 | tyrosine kinase with immunoglobulin like and EGF like domains 1 | ENSG00000066056 | 2,19E-08 | 2,93 |
| ACLY | ATP citrate lyase | ENSG00000131473 | 1,56E-09 | 2,92 |
| PARD6A | par-6 family cell polarity regulator alpha | ENSG00000102981 | 1,52E-10 | 2,89 |
| IL36B | interleukin 36 beta | ENSG00000136696 | 6,17E-11 | 2,86 |
| ITGAL | integrin subunit alpha L | ENSG00000005844 | 1,48E-09 | 2,75 |
| HTRA1 | HtrA serine peptidase 1 | ENSG00000166033 | 1,78E-08 | 2,69 |

Abbreviations: False discovery rate corrected p-value (FDR); Median fold change (FC).

**Table 4.**
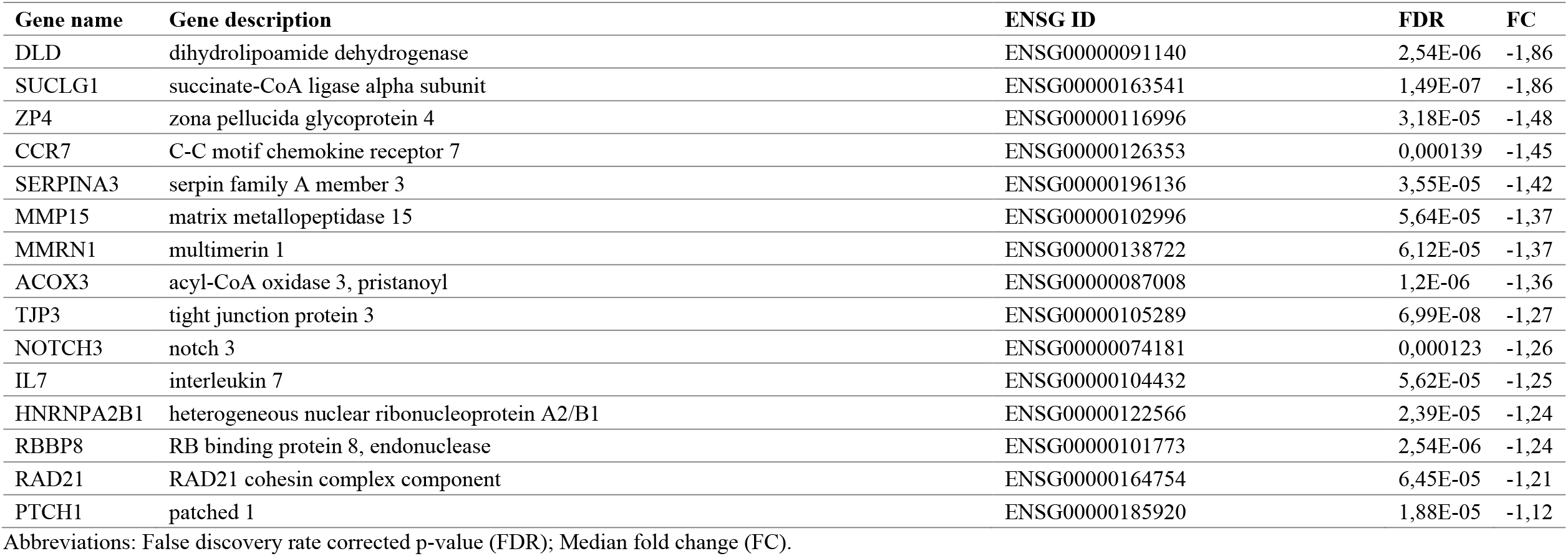
Proteins with higher levels in cluster 1 compared to the other clusters.

## Discussion

Applying an unsupervised analysis approach on plasma proteomic data from women of the KARMA breast cancer risk cohort, we identified a subset of individuals with more previous use of MHT and a greater proportion of breast cancers. The women in this cluster were also older and had a larger MD area relative to their age. Characterization of circulating proteins driving the cluster found an lower levels of proteins involved in cell adhesion and immunoregulation, and a higher levels of proteins associated with DNA integrity, cell fate, metabolism and the female reproductive system.

Data-driven archetypal analysis was used as an unsupervised approach to identify proteomic-based clusters in the data, which were linked to phenotypic or genotypic traits in a population. This enabled the identification of associations between clusters of women with similar plasma profiles and risk factors for breast cancer. By clustering the participants based on the proteomics data, we found strong associations with previous use of MHT, where 79% of participants in cluster 1 were previous users.

In the same cluster we also found an overrepresentation of breast cancers, with 58% being cases compared to 28-35% in the other clusters. This confirms previous knowledge that the use of MHT is associated with an increased 5-year risk of breast cancer among postmenopausal women [36]. Of note, all cases in cluster 1 had previously been treated with MHT, while this was only true for half of the cases in other clusters. The proteomic signature of cluster 1 associated with MHT usage, however, this was not driven by current use of MHT. This suggested that previous use of MHT left a mark in the circulating proteome of these women that could be detected even years after discontinuing the treatment. Individuals in cluster 1 also had a greater mammographic density relative to their age which is a known risk factor for breast cancer. Interestingly, MHT usage is known to be associated with higher mammographic density in postmenopausal women [37-41]. However, to our current knowledge, no longitudinal studies have been performed to investigate potential long-term effects of MHT on density. Our results suggest that such studies may be warranted. It is therefore not clear if the increased relative density observed in cluster 1 is due to the previous MHT use or other factors. Interestingly, statin use was not seen as a major driver of the protein profiles despite their known effects on the plasma proteome supporting that the observed effect of MHT is specific for this class of drugs. Additionally, no effect of genetic risk was observed, however, this could be due to too low sample size.

The shortlisted set of proteins targets with differential abundance in cluster 1 compared to the other clusters were related to DNA repair/integrity, cell fate/replication, mammographic density, and the female reproductive system, thus supporting their putative roles in development of breast cancer or mediation of risk factors.

We found that individuals in cluster 1 had lower levels of circulating proteins regulating DNA repair/integrity (RBBP8, RAD21) and cell fate/replication (NOTCH3, TJP3, HNRNPA2) thus indicating a role in cancer development. In concordance with this, we found RBBP8, TJP3 and HNRNPA2 to be significantly, negatively associated with mammographic density. Individuals in cluster 1 had higher circulating levels of proteins that may be linked to mammographic breast density and the accompanying mechanical stiffness. These proteins included the cell junction and adhesion molecules CLDN15, ITGB7, F11R and its receptor ITGAL, which are potentially involved in sensing of stiffness in the breast tissue and activation of cellular downstream signaling pathways to maintain tissue homeostasis [42-46]. In line with this, we also found these proteins to be positively associated with mammographic density, though the associations were not significant. Reassuringly, we have previously found, in two separate data sets, positive associations between mammographic density and F11R [20]. In fact, F11R has been widely described in cancer development and progression and the expression of F11R correlates with poor breast cancer prognosis [47, 48]. Our current findings validate our previous results and supports our hypothesis that F11R plays a role in regulating mammographic density and breast tissue composition.

The levels of the proteins ZP4 and PTCH1 were found to be lower in cluster 1. Additionally, across clusters, the two proteins were decreased for cases compared to controls and in MHT treated compared to untreated women. Both proteins are expressed in female tissues and we found both proteins to be negatively associated with mammographic density. Interestingly, these were the only two cluster-1-specific proteins that were also significantly associated with MHT use. We therefore hypothesize that MHT might negatively affect the expression in female tissues and thereby affect the plasma abundance of these proteins. ZP4 was selected for inclusion in this study due to its role in extracellular matrix (SBA1). It is primarily expressed by the ovary and placenta, but also other tissues [22, 49]. ZP4 is part of the extracellular matrix surrounding oocytes and has been linked to the fertilization processes [50, 51]. The protein PTCH1 was included in this study as it has previously been linked to cancer (SBA3). As a protein found on the cell surface and the Golgi apparatus, it functions as a tumor suppressor, and mutations of the *PTCH1* gene have been associated with poor prognosis and increased recurrence of breast cancer [52]. PTCH1 is expressed more widely than ZP4, but is among many tissues, expressed in female tissues, especially the cervix and endometrium [22, 49]. The two proteins, ZP4 and PTCH1, could therefore potentially represent an unknown link between MHT usage, female tissues and mammographic breast density all leading to increased risk of breast cancer.

Apart from cluster 1, the assigned members of the remaining clusters showed high interchangeability among another when the data was perturbed. They should therefore only be interpreted with caution [28]. Clearer definition criteria for these clusters could possibly be achieved by applying stricter inclusion cut-offs where any unassigned participants are further pooled into “in-between” groups corresponding to individuals who do not reliably belong to single clusters. This possibility is also one of the strengths of archetype analysis over more traditional and static clustering methods. The non-binary cluster membership offers a greater flexibility to reflect the extent of the diverse processes of human biology. However, such investigations go beyond the scope of this work. Consequently, we chose to focus on the clearest difference observed between women in the most stable cluster 1 and the remaining cohort.

Using a traditional approach to compare cases and controls by their proteomic profiles, we found no targets to be statistically significant. This is in line with previous literature reporting few or no protein biomarkers for overall early detection of breast cancer [2-8]. Likely, this reflects the already early detection possible by mammographic screening, the complex etiology and heterogeneity of the disease, and that effects from a multi organ system contribute to the granularity in the circulating plasma proteome. Most previous attempts have identified putative subtype specific markers with, at best, limited performance in replication and validation efforts. Herein, we did also not detect any significant subtype-specific profiles of circulating proteins deemed useful for early detection.

Weaknesses in our study can be seen in the low number of breast cancer cases available from prospective studies. Other weaknesses can relate to an initial sampling of participants based on a classical case-control design with two matched controls for each breast cancer case. As the case-control analyses provided limited insights, we proceeded with a data-driven, thus hypothesis-generating strategy. Therefore, the cohort of women included in this study were enriched for breast cancer cases compared to the general population. However, this enrichment of cases increased the chances of observing effects related to risk factors and case-control status where much larger numbers of participants would otherwise have been needed. Furthermore, we used plasma to identify proteomic signatures associated with breast cancer risk factors and early detection. As previously discussed [20], it remains to be ascertained how well circulating protein concentrations reflect the changes in the protein expression of the breast tissue. However, as we have shown here, it seems that several systemic processes contribute to the physiological changes occurring in breast cancer patients, and plasma provides a window into processes occurring in multiple tissues in one go. Nevertheless, the identified epithelial and stromal cell-specific proteins support protein leakage or shedding into blood, and that an elevated turnaround of proteins in breast tissue can lead to the detection of these targets in the circulation. Although we are using the very-well characterized hence comprehensive KARMA cohort, information on tumor characteristics and risk factors was missing for some participants. In particular, data specific to MHT subtypes, dosage, and duration of the treatment, as well as some information on tumor characteristics, was missing. Exposure data in KARMA is self-reported, which may result in measurement bias. However, exposure data, mammograms and blood samples were collected at the same time at KARMA study entry, and it is not likely that the participants knew about their mammographic density at the time of answering the questionnaire. Besides, a non-differential misclassification of exposures would dilute, not strengthen, the reported associations. Additionally, questionnaire data on drug usage was supplemented with data from the Swedish drug prescription registry. Given the expected heterogeneity of the molecular phenotypes, a lack in power may have further weakened the statistical significance of our findings. Our observations further prompt validation in an independent cohort and dataset of a comparable design and depth.

Strengths of our study reside in the utilized exploratory affinity-based proteomic assay. It provides novel opportunities for high-throughput screening for circulating proteins associated to risk factors, indicative for disease development in selected phenotypes. The experimental design allows combining different protein assays into one multiplexed approach and it is attractive due to it consumption of only minimal sample volumes. The method reports relative protein quantities in plasma that allow a comparative analysis across different samples. Strengths also include the centralized and standardized collection of high-quality blood samples, which is also evident from the fact that we observed no systematic differences at the protein level between sampling centers. Additionally, the centrally managed questionnaire data and mammograms obtained from all KARMA cohort participants prior to diagnosis, as well as the quantitative assessment of mammographic density by STRATUS [53] are strengths of this study.

## Conclusion

Our findings suggest that use of MHT may leave long-lasting fingerprints in the circulating proteome. Effects of the treatment could be detected in the proteome even years after discontinuation and were especially apparent for proteins associated with mammographic density and breast tissue composition, tumor development and progression, and the female reproductive system. Like previous studies, we did not identify any independent markers of early detection of breast cancer from plasma proteins. Instead, we identified circulating proteins associated with previous MHT use, connecting to a higher frequency of women with breast tumors, greater age and relatively greater mammographic density. These findings provide novel biological insights to putative pathological processes associated with MHT usage and breast cancer risk. Collectively, this suggests that rather than looking for biomarkers secreted by a developing tumor for early breast cancer detection, proteomic characterization of plasma might be more successfully aimed at identification of biomarkers that modify or explain the effects of known risk factors, and that unsupervised analysis approaches may aid in this endeavor by providing novel hypotheses. Our findings need to be further validated, in both plasma and in breast tissue, but they support the notion that further integration of health and treatment trajectories need to be considered when judging some of the molecular phenotypes of a disease.

## Declarations

### Ethics approval and consent to participate

All participants signed informed consent forms before joining the KARMA study, and the ethical review board of Karolinska Institutet approved the study (2010/958-31/1).

### Consent for publication

All authors approved of the manuscript and consented to its publication.

### Availability of data and material

The datasets used and/or analyzed during the present study are available from the corresponding author upon reasonable request. Code developed for performing the analyses available at: https://github.com/Schwenk-Lab/karma_breast_cancer

### Author contributions

JMS, MG, SB and CET conceived and designed the study. PH and JMS supervised the study. SB generated plasma proteomics data. LD performed the statistical and data driven analyses. CET and LD lead the interpretation of the data with support from all co-authors. CET and MG drafted the manuscript with support from LD and JMS. SB, YC, AM, MU, KC and PH critically reviewed the manuscript. All authors read and approved the final version of the manuscript.

## Acknowledgements

We thank all the participants in the KARMA study, the study personnel for their devoted work during data collection Also, we thank the everyone from the Human Protein Atlas for their efforts. We thank Mun-Gwan Hong and Tea Dodig-Crnkovic for the fruitful discussions, and the Translational Plasma Profiling facility at SciLifeLab for the support in generating the data for this project. Figure 1 was created with BioRender.com.

